# PhiDsc: Protein functional mutation Identification by 3D Structure Comparison

**DOI:** 10.1101/2022.05.18.492407

**Authors:** Mohamad Hussein Hoballa, Changiz Eslahchi

**Affiliations:** Department of Computer Science, Shahid Beheshti University, Evin, Tehran, 1983963113 Iran

## Abstract

Selective pressures that trigger cancer formation and progression shape the mutational landscape of somatic mutations in cancer. Given the limits within which cells are regulated, a growing tumor has access to only a finite number of pathways that it can alter. As a result, tumors arising from different cells of origin often harbor identical genetic alterations. Recent expansive sequencing efforts have identified recurrent hotspot mutated residues in individual genes. Here, we introduce PhiDsc, a novel statistical method developed based on the hypothesis that, functional mutations in a recurrently aberrant gene family can guide the identification of mutated residues in the family’s individual genes, with potential functional relevance. PhiDsc combines 3D structural alignment of related proteins with recurrence data for their mutated residues, to calculate the probability of randomness of the proposed mutation. The application of this approach to the RAS and RHO protein families returned known mutational hotspots as well as previously unrecognized mutated residues with potentially altering effect on protein stability and function. These mutations were located in, or in proximity to, active domains and were indicated as protein-altering according to six *in silico* predictors. PhiDsc is freely available at https://github.com/hobzy987/PhiDSC-DALI.

## INTRODUCTION

Cancer development starts with the acquisition of genomic alterations and chromosomal abnormalities that arise from uncorrected errors during DNA replication or repair or due to exposure to mutagens (1). Some alterations may further the accumulation of somatic mutations (2) and play a mechanistic role in malignant transformation. These “driver mutations” are postulated to provide advantage to and promote cancer hallmarks in the subpopulation of cells that harbor them (3). The number of driver mutations varies between cancer types, averaging four per tumor (4). Most remaining somatic alterations, termed “passenger mutations,” may confer little to no functional impact (5). However, distinguishing the handful of driver mutations from the vast background of passenger mutations in a tumor has remained a challenge in cancer genomics.

Frequently altered nucleotides in the genes that are implicated in tumor development and progression are known as mutational hotspots (6). The number of candidate hotspot mutations of unknown functional significance has increased recently–especially due to the completion of large-scale sequencing efforts such as The Cancer Genome Atlas (TCGA) (7), International Cancer Genome Consortium (ICGC) (8), and Project GENIE (9). Many platforms are used to visualize and organize these data like BioMuta (10) and cBioPortal(11, 12) allowing to download and analyze large-scale cancer genomics datasets. Most of these frequently detected mutations are within exons, or the coding regions of the proteins, and their function is ascertained by directly examining their impact on the encoded protein or predicted through application of *in silico* bioinformatic approaches (13, 14).

The statistical reoccurrence of mutations in tumors has been used as an indicator of their functional impact, based on the assumption that infrequent alterations detected in tumors are likely non-functional, passenger events (15). However, it has been shown that passenger mutations are not randomly distributed along the cancer genomes (16). Rather, they are enriched in nucleotide sequence contexts that are shaped by specific active mutational processes in a tumor (17, 18). In contrast, driver mutations are postulated to occur in genomic positions whose distribution depends not only on the local nucleotide context, but also on the location of functionally relevant residues along the protein sequence (19, 20). Relying on recurrence alone to identify functional mutations, may also be confounded by underlying mutational processes that target specific genomic contexts, resulting in often-mutated residues that do not drive tumor progression (21).

In this context, numerous methods are presently being used to identify hotspot and driver mutations, based on the frequency of mutations detected in a gene across a set of tumor samples (e.g., MutSig (22) and MuSiC (23)). Recognizing mutational hotspot in infrequently altered genes can also be refined by including protein-level annotation by local-positional clustering (24), or the inclusion of phosphorylation sites (25) and information from paralogous protein domains (26). Protein-level annotation, such as local-positional clustering, phosphorylation sites, and paralogous protein domain (27) as well as 3D protein structures are used to identify functional mutations in infrequently mutated genes.

Using a variety of approaches that take into account diverse aspects of protein structures and types, functional mutations can be predicted across protein sequences and structures. Some techniques, such as 3DHotspots (28), Hotspot3D (29), Mutation3D (30), and Signatures of Cancer Mutation Hotspots in Protein Kinases (31) use the 3D structure of protein, while others utilize 3D reconstruction of protein networks to provide a better understanding of genetic abnormalities (32). On the other hand, methods like PinSnps (33), StructMAn (34), Hot-MAPS (35) and SpacePAC (36), as well as SAAMBE-3D(37), use protein-protein interactions enriched with somatic cancer mutations (38) to understand the effect of a mutation not only on the function of the same protein but also on the signal transduction and activating cascade proteins. Methods based on individual protein structures or the 3D reconstruction of protein networks have improved the identification of mutational clusters in tumors (39) and have elucidated functional consequences (folding free energy and stability of protein monomers (40)) of protein-altering mutations, other methods take into consideration the local DNA sequence context for the analysis of cancer context-dependent mutations like MutaGene(41). Although it is difficult to categorize methods based on their input parameters (some require sequences while others may need structures as well), in all cases, the output determines whether a proposed mutation has occurred at a hotspot residue. However, a few limitations remain: First, focusing on the mutation frequency across tumor samples increases the risk of missing portions of rare hotspot mutations with low frequency; second, concentrating solely on driver genes fails to distinguish between individual driver mutations within altered genes and passenger mutations within the same gene; and third, analyzing protein sequences without a larger context misses the effect of mutations on the conformational structure and functional sites of the protein.

To address these issues, we introduce PhiDsc. Its development is based on the hypothesis that oncogenic mutations in a target protein can be identified by analyzing its three-dimensional structural similarity, protein folding information, and mutational recurrence within its gene family. We demonstrate that PhiDsc can identify candidate functional mutations, caused on altered protein position, by comparing the three-dimensional structures of related human wild-type proteins and assessing repeatedly altered residues in the protein family. PhiDsc combines the two approaches by relying on the concept of hotspot mutations in functional regions and classifying protein families based on their domains and active sites. Thus, by comparing the three-dimensional structures of similar domains within a protein family, PhiDsc maps known functional mutations in extensively studied proteins to those in the family that receive less interest.

## RESULTS

PhiDsc is applied to HRAS from the RAS (59) subfamily and RhoA from the RHO (60) subfamily of proteins.

### HRAS

The family group of HRAS was A(HRAS) = {DIRAS1, DIRAS2, GEM, KRAS, NRAS, RAP1A, RAP1B, RAP2A, RASL12, REM1, REM2, RERG, RRAD, RRAS, RRAS2}. Dali aligned 98% of HRAS residues to residues of each member of the family **(Table 1)** highlighting strong structural similarity between the target protein and its respective protein families. (Supplementary files HRAS alignment). As a result, PhiDsc scored 168 of 189 HRAS residues (89%) and predicted 13 residues as functional mutation (**Table 2**) all of which passed cross-validation evaluation (**Figure 1**) and were consistently projected to be effective and protein-modifying by six independent algorithms.

**Table 1.**
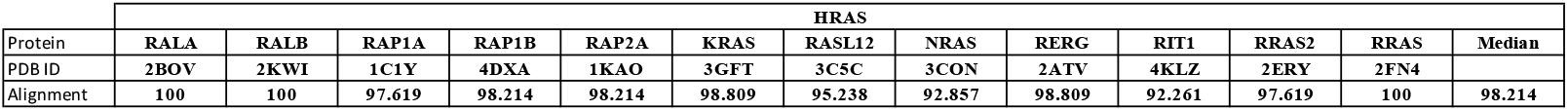
indicates the percentage of structural alignment between each protein (HRAS) and its protein family member.

**Table 2.** Candidate functional mutations for HRAS proposed by PhiDsc. Residue positions sorted by their PhiDsc score p-value along with predicted interacting residues from the RIN analysis are shown. COSMIC mutation reference or dbSNP polymorphism ID are

| HRAS |  |  |  |  |  |  |  |  |  |  |  |
| --- | --- | --- | --- | --- | --- | --- | --- | --- | --- | --- | --- |
| Residue Nu | P-value | Interacting Residue |  |  |  |  |  |  |  |  | mut ref Nu |
| 12 | 2.66E-07 | 11 | 16 |  |  |  |  |  |  |  | COSM483 |
| 74 | 1.72E-06 | 5 | 70 | 71 | 73 | 75 |  |  |  |  | COSM5991570 |
| 13 | 2.57E-06 | 117 |  |  |  |  |  |  |  |  | COSM486 |
| 93 | 6.18E-06 | 81 | 82 | 90 | 91 | 113 | 137 |  |  |  | COSM9497546 |
| 91 | 8.72E-06 | 87 | 88 | 90 | 93 | 95 |  |  |  |  | COSM6476473 |
| 22 | 1.38E-05 | 18 | 19 | 20 | 32 | 26 | 28 | 146 | 149 | 152 | COSM6923245 |
| 96 | 1.54E-05 | 9 | 10 | 11 | 92 | 93 | 97 | 98 | 99 | 100 | RS889495169 |
| 117 | 1.85E-05 | 13 | 14 | 83 | 84 | 116 | 119 | 120 | 144 |  | CSOM304967 |
| 31 | 3.84E-05 | 30 | 33 |  |  |  |  |  |  |  | COSM6915342 |
| 40 | 4.20E-05 | 20 | 24 | 32 | 38 | 39 | 54 | 55 | 57 |  | RS763920334 |
| 155 | 5.08E-05 | 79 | 144 | 151 | 152 | 153 | 159 |  |  |  | COSM9515051 |
| 148 | 5.23E-05 | 119 | 145 | 150 |  |  |  |  |  |  | COSM6903495 |
| 38 | 5.93E-05 | 39 | 40 | 57 |  |  |  |  |  |  | RS750680771 |

**Figure 1.**
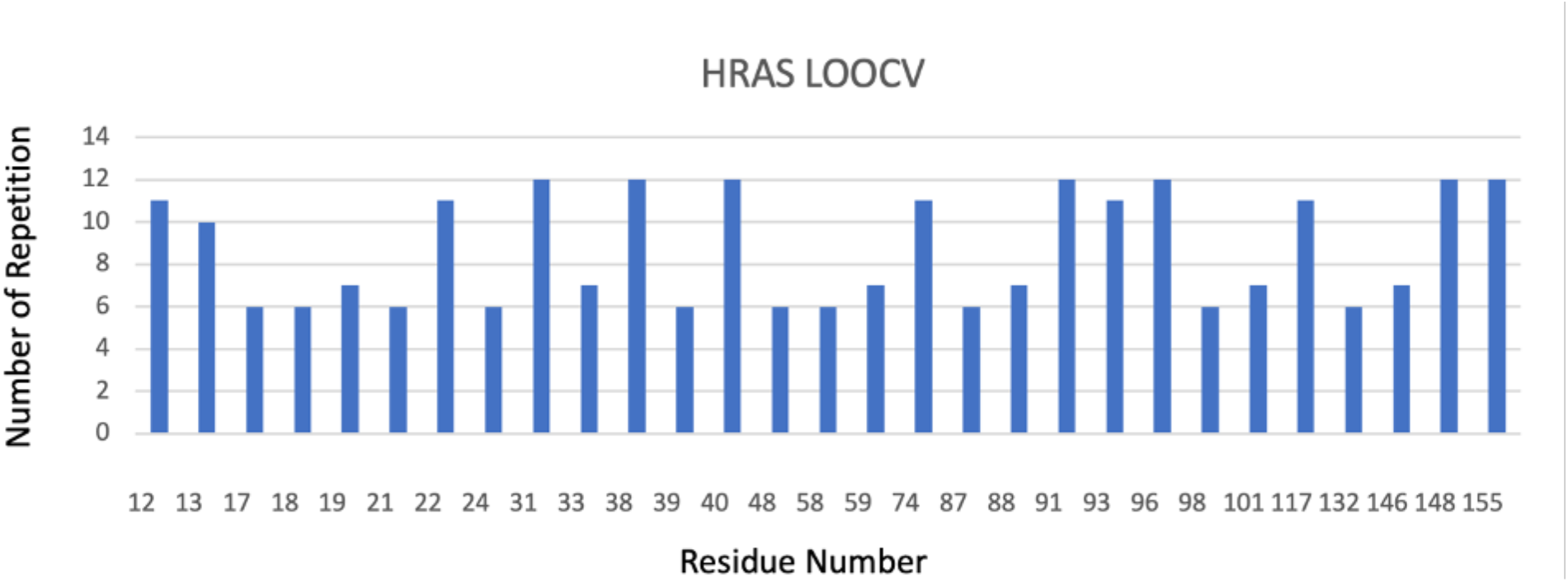
shows the LOOCV for the two proteins HRAS, in all the iterations of the system the number of repeated times for each residue is shown, (>80%), which indicates that the results obtained by the system are robust since the original results are obtained in all the LOOCV iteration

RIN is generated using the HRAS structure (RCSB database ID 4Q21, with 168 residues). Thirteen candidate functional mutations shared 58 neighboring residues located in the functional domains of the protein (G boxes, Switches I and II, GDI and GEF interaction sites, GTP/MG2+ binding domain). Moreover, 25 of these 58 residues were seen mutated in human tumors according to the cBioPortal(11, 12) database a distinct dataset form BioMuta.

Top-four PhiDsc predictions in HRAS were residues 12, 13, 74, and 93, which are known to be key functionals and often mutated in various cancer types (61). The domain comprising residues 12 and 13 is involved in Guanine Nucleotide Dissociation Inhibitor (GDI) interaction as well as interaction with GTP/Mg2+ (62), and is mostly detected in tumors such as bladder cancer (63), thyroid cancer(64), and other diseases such as Costello syndrome (61) and Schimmelpenning-Feuerstein-Mims syndrome (63, 65). Mutations in residue 74 are seen in endometrioid cancer and sebaceous carcinoma, while those in residue 93, have been discovered in only a small percentage of prostate cancer samples (66). According to Ensemble Learning Approach for Stability Prediction of Interface and Core mutations (ELSPIC) (67), residue 93 is localized in the protein’s core, suggesting that it has a direct effect on the protein’s shape and function.

Although 3 of 13 candidate functional mutations in HRAS were not located in any protein domains, they were found near the intersection of exons 3 and 4 at residue 97. Finally, residue 96 has been identified as a phosphorylation site, the other residues as showen in (**Figure 2**) were located in functional protein domains.

**Figure 2.**
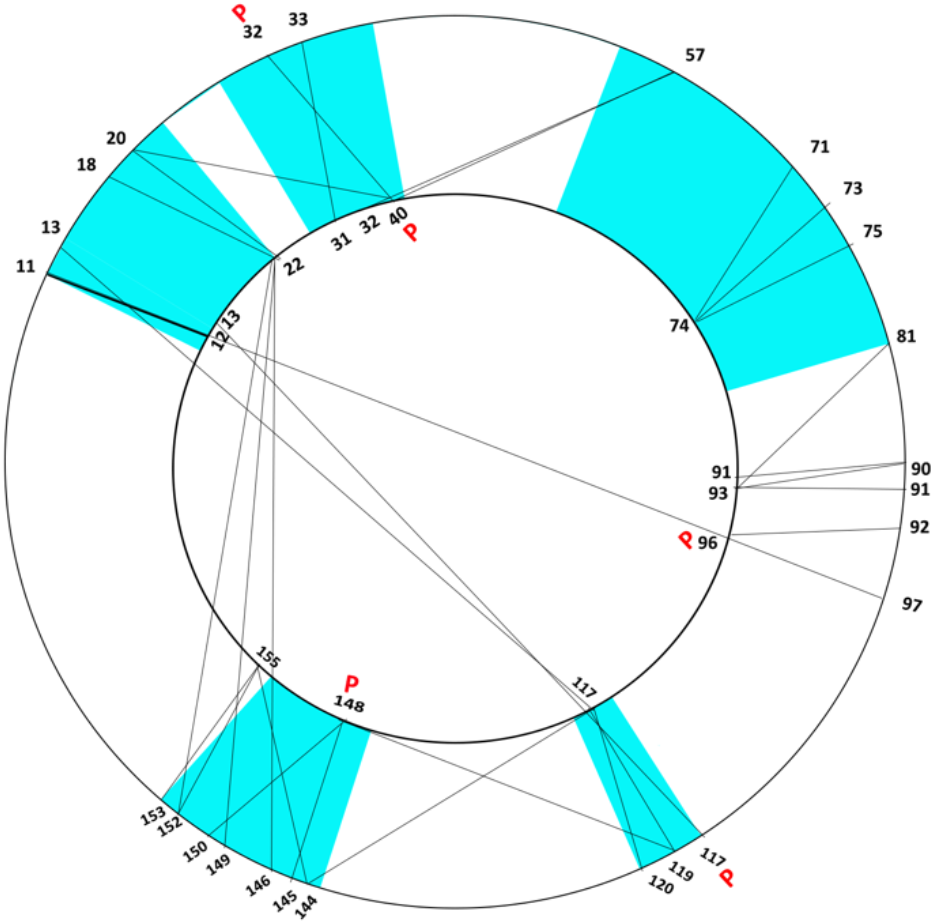
depicts the inner circle’s candidate functional mutations and the outer circle’s interacting residues. According to thecanSAR BLACK system (60), The blue areas represent HRAS functional regions, while the lines linking the inner circle (candidate functio functional mutation) to the outer circle (interacting residues) represent residue interactions. This figure displays only the HRAS residues that are mutated in cBioPortal.

### RhoA

RhoA, a member of the RHO (60) subfamily of proteins with A(RhoA) = {RHOB, RHOC, RHOD, RHOQ, RHOU, RND1, RND3, RAC1, RAC2, RAC3, CDC42}.

The RCSB database is used to retrieve 3D structure files for each member (if found in PDB) of A(RhoA). The final list of PDB structures are shown in **Table 3**. The Dali server is then used to perform a pairwise structural comparison between the input protein and each member of its family. 97% of RhoA residues were aligned with the residues of each family member in the generated alignments. The existence of strong structural similarities between target proteins and their respective protein families supports these results (Supplementary file “RhoA alignment”).

As an outcome, 179 out of 193 residues were scored for RhoA.

**Table 3.**
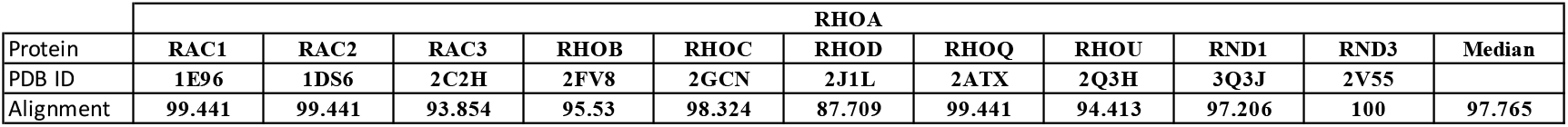
shows the percentage of structural alignment of each protein (RhoA) with its corresponding protein family member.

The P-value of PhiDsc statistics is generated for all target protein residues in the final phase. Eight candidate functional mutations for RhoA were obtained. **Table 4** illustrates the RhoA protein candidate functional mutations introduced by the PhiDsc procedure. The eight candidates passed cross validation (**see Figure 3**) and were consistently predicted to be effective and protein-modifying by six separate algorithms. Despite the fact that no evidence of a mutation in residue 29 of RhoA was detected in any cancer mutation databases, all six techniques predicted that this mutation would alter RhoA’s functional activity.

**Table 4.** lists all candidate functional mutations for RhoA proposed by the PhiDsc approach. The table shows the residue position number (P) in the first column, sorted by their P-value in the second column, the interacting residues of each candidate functional mutation in the third column, the “COSM” letters of the mutations indicate that these mutations were annotated in the cosmic database as tumor-related mutations, while the “rs” letters of the mutations indicate that these mutations were annotated in the Dpsnp database.

| RHOA |  |  |  |  |  |  |  |  |  |  |
| --- | --- | --- | --- | --- | --- | --- | --- | --- | --- | --- |
| Residue Number | P-value | interacting residue |  |  |  |  |  |  |  | mutation ref NU |
| 111 | 3.07904E-05 | 78 | 79 | 80 | 109 | 110 | 177 |  |  | COSM2849881 |
| 34 | 4.86819E-05 | 35 |  |  |  |  |  |  |  | COSM2849895 |
| 139 | 9.956E-05 | 84 | 86 | 89 | 92 | 122 | 139 | 140 | 143 | COSM2849897 |
| 168 | 0.000147858 | 170 | 171 | 172 |  |  |  |  |  | COSM7114068 |
| 110 | 0.000209526 | 77 | 78 | 79 | 80 | 107 | 108 | 11 |  | RS368767616 |
| 29 | 0.000224094 | 23 | 27 | 28 | 29 | 31 |  |  |  | NO |
| 172 | 0.000300752 | 46 | 48 | 168 | 169 | 172 | 174 | 175 | 176 | COSM1309264 |
| 127 | 0.000484266 | 87 | 121 | 124 | 125 | 127 | 129 | 130 | 131 | MU85445108 |

**Figure 3.**
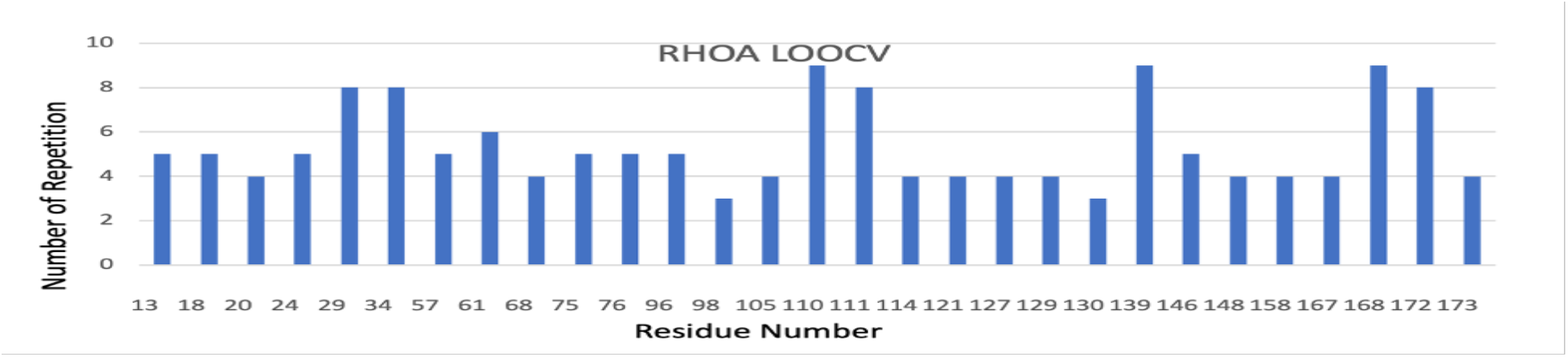
shows the LOOCV for the protein RhoA; the number of repeated times for each residue is presented in all iterations of the system, indicating that the system’s results are resilient because the original results are obtained in all LOOCV iterations.

The RIN for RhoA is constructed using 1OW3 obtained from the RCSB database. The 8 potential functional mutations have 42 neighbors, 18 of which had previously been identified as occurring mutations in the cBioPortal database (11, 12) (**see Table 3/interacting residues**). The neighbors of potential functional mutations are related to PPI functionals, according to RINalyzer data. These neighbors are also located in RhoA protein domains associated to GAP, GEF, and GDI interaction and phosphorylation sites, including position 127—showing that this residue is significant in RhoA’s functional activity (**see Figure 4**).

**Figure 4.**
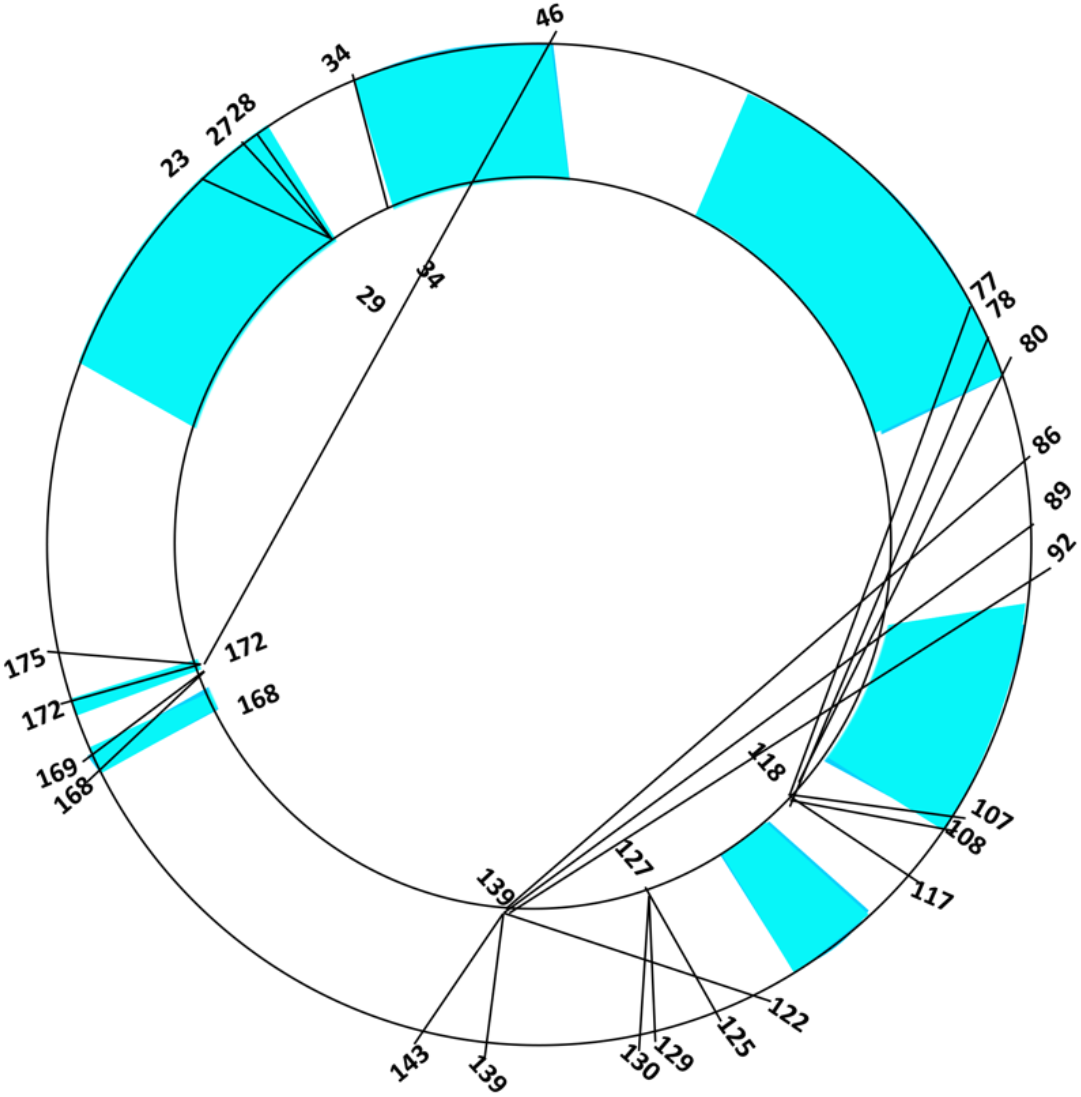
showes the inner circle’s candidate functional mutations and the outer circle’s interacting residues. According to thecanSAR BLACK system (60), The blue areas represent RhoA functional regions, while the lines linking the inner circle (candidate functional mutation) to the outer circle (interacting residues) represent residue interactions. This figure displays only the HRAS residues that are mutated in cBioPortal.

In cancer samples, four high-scoring RhoA residues (34, 139, 111, and 168) were observed (**see Table 4)**. Residue 34 is near the core area and the GAP interaction site, as per RhoA’s 3D structure. A mutation at this location improves the affinity for ARHGAP;1, a GAP protein that plays a vital role in RhoA activation, according to data from ELASPIC (67) and COSMIC. According to COSMIC, mutation 139 of RhoA was observed in one sample of non-small cell lung carcinoma and as a silent mutation in two samples of cervix and stomach cancer— where it was not a functional mutation in the latter two samples. Meanwhile, residue 111 has been seen in one sample of stomach cancer patients (7). Mutation in residue 168 boosts the affinity for the CTRO protein, which regulates cytokinesis by generating a contractile ring. It was also found to interact with KAPCA, a gene associated with breast and ovarian cancer (68). The mutation of residue 168 also impacted PKN1 and PKN2 interaction with RhoA—two proteins that contribute to prostate cancer and play a crucial role in cell migration and proliferation (69, 70).

## DISCUSSION

In this paper, we looked at proteins that are similar and have been classified into families in uniportkb. In terms of sequence, structure, and function, these proteins are very similar. As a result, we assume that the frequent mutations associated with the same cancer phenotype on the same domain share these domains and mutations within the family. As a result, the introduced algorithm employs scores to determine whether these mutations are statistically significant as functional alterations in areas common in families. To test and validate the approach, domains from two well-known protein families (HRAS and RhoA) that are known to be involved in cancer are used.

As a result, we present PhiDsc, a novel method for detecting functional mutations in proteins. To link mutation residues to specific biological functional domains of proteins, we took into account a mutation’s position in the protein’s 3D structure (71), as well as the frequency of its reoccurrence in human tumors (72). Finally, we combined these characteristics with known functional hotspot mutations aggregated among paralogous proteins in the same family or with similar domains (73), and we used Bonferroni restriction to further narrow the range of predictions in order to reduce false positives..

We evaluated PhiDsc using the HRAS and RhoA proteins (71, 72). HRAS is a GTPase protein in the RAS subfamily that controls many cellular mechanisms including 84 pathways according to KEGG Pathway. The most mutated residues in HRAS are 12, 13, and 61, which are related to different subsets in cancer (73), and the tumorigenic effect of HRAS is related to the protein’s permanent activation. RhoA is a RHO subfamily signaling G protein that regulates numerous cellular mechanisms associated with 43 pathways related to cellular processes as seen in KEGG Pathway. The most frequently mutated residues in this protein, 17 and 42, have been observed in various types of cancer (74), and similarly, the oncogenic effect of RhoA is exerted by its constant activation of the protein.

With the exception of one candidate residue in RhoA, all residues predicted by PhiDsc were found to be mutated in cancer samples, as well as in other diseases such as Costello syndrome, which is linked to germline HRAS mutations (75). Although certain candidate functional mutations were not previously identified as hotspot mutations and had a low mutated frequency in cancer mutation datasets (rare), using CanSar balck (60), we demonstrated that they were located in active functional domains of proteins or had a wide network of interactions with functional residues. Noteworthy, the Biomuta database was initially used; however, by the final step, some of the candidate functional hotspots that were not found in Biomuta had been presented in tumor samples in other datasets such as COSMIC, cBioPortal, and Dbsnp. With the exception of RhoA residue 29, all were identified as rare mutated residues, and, thus, they were not previously mentioned as a hotspots, indicating that PhiDsc improves and optimizes the detection of low frequency functional mutations. while, residue 29 of RhoA had no mutational recorde in COSMIC (46) or Dbsnp (76) databases, mutation analysis software MutaGene (41) ranked RhoA residue 29 as a highly mutable position, and the projected effect by six different software packages at that position predicts a potential oncogenic effect. It is notable that the difference between COSMIC and Dbsnp lies in the curation method used to classify any given mutation as an SNP.

Despite the fact that these methods use different concepts to infer the stabilizing effect of point mutations (as discussed in the results section), they all suggest that PhiDsc’s predictions alter protein structure and function. The precise impact of unknown mutations necessitates additional experimental verification.

When DALI was used instead of TM-Align, better results were obtained in PhiDsc with known functional mutations. These findings suggest that different 3D alignment approaches may alter predicting hotspot mutations in different types of proteins. As a result, the PhiDsc package’s predictions should improve as the mode of alignment used improves.

Some previously designated hotspots of HRAS and RhoA in cancer, like for HRAS out of 12 (residues 12, 13 and 117) and for RhoA out of 11 (residue 34) were returned by PhiDsc. When the results of the Dali and Tm-Align alignment (supplementary files (HRAS, RHOA) Tm-Align) methods were compared, the results of the Tm-Alignment method predicted fewer well-known driver mutations than the results of the Dali method. This suggests that a different alignment choice could result in some differences in predictions.

Although the two example proteins selected for validation are oncogenic, PhiDsc is not restricted to oncogenes and can be utilized to identify functional mutations in tumor suppressor genes or any other type of Protein if the family has a sufficient number of members and the mutation profile data is adequate and consistent.

The lack of a 3D structure of the protein and small protein families, which limit the number of members in the family, are two limitations of this method. A future update to the tool will include the ability to align functional domains of proteins rather than the entire protein, as well as the use of the protein’s predicted 3D structure in the alignment comparison.

## MATERIALS AND METHODS

### PhiDsc Algorithm

PhiDsc uses a six-step method that is centered on a protein **P** with **m** amino acid residues and a known three-dimensional structure. Briefly, a list of proteins is defined, denoted by the set **A(P)**, by identifying all members of P’s protein family from UniProtKB (42) and selecting all human proteins with 3D structures from the Protein Data Bank (PDB) (43). Next, the 3D structures of the proteins members in A(P) are aligned to the 3D structure of P. The results are presented by a matrix, **E(P)**. Then, using the BIOMUTA V4 and 3Dhotspot database (44), the mutational information of each protein of A(P) is identified, in order to score each residue of P and calculate an associated probability. Finally, these are analyzed to identify potential candidate functional mutations in P. Each step is described in detail in what follows.

Step 1: Define the protein list A(P). The UniProtKB database (42) is used to identify members of a given protein’s protein family, while the RCSB Protein Data Bank (PDB) (43) is used to determine their three-dimensional structure. The PDB contains the structures of wild-type and mutated proteins. For the alignment step, either the full-length sequence of the wild-type protein or the least mutated form (maximum one mutation) of the same length is used; the final list is denoted by *A*(*P*) = {*p*_1_,*P*_2_,*P*_3_… *P*_n_}.
Step 2: Align 3D structures. Dali, a pairwise comparison server for protein structures, is used to align protein structures (http://ekhidna2.biocenter.helsinki.fi/dali/)(45). TM-Align “another alignment method” is also included in PhiDsc with its default parameters.
Step 3: Define matrix E(P). 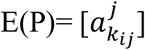 has n columns (number of proteins) and m rows (number of amino acids in protein P), in which 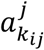 denotes the type of amino acid in the sequence of protein j that is aligned to the *i^th^*amino acid in protein P; k_ij_ denotes the position number of amino acid in the sequence Pj that is aligned to the *i^th^* amino acid of protein P.
Step 4: Identify mutational information of each protein in A(P). Residues for all protein family members are annotated with mutational and hotspot information using BioMuta (version 4, (10)) and 3Dhotspots (39). BioMuta is a database of curated cancer-associated single-nucleotide variations derived from COSMIC (46), ClinVar (47), CIVIC(48), and UniProtKB(42) and actively curated from publications and automated analysis of publicly available databases such as TCGA(7)and ICGC(8). 3Dhotspots is a dataset of statistically significant mutations clustered in three-dimensional protein structures found in cancer. The data set contains mutational positions referred to as hotspot mutations.
Step 5: Score residues. A grade is assigned to each amino acid of A(P) members based on the mutational information for that amino acid (P). Let 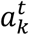 be the kth amino acids of protein *P_t_*. Define:

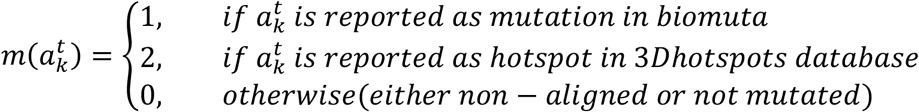 Let the *i^th^* row of the matrix E(P) be 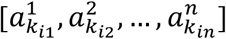, 1 ≤ *i* ≤ *m*. The following score is assigned to *i^th^* amino acids of P:

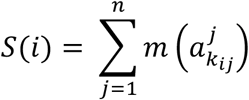 To calculate the statistical significance of the obtained scores *S*(*i*) at each position (row in the matrix E(P)), we calculate the probability related to this score. Let protein *P_t_* have *m_t_* amino acids of which *l_t_* are mutated in biomuta. Define:

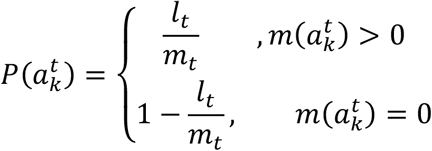 To distinguish non-mutated from the non-aligned residues (both with score 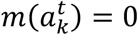), and because the event under investigation is the occurrence of functional mutations that are coded in the alignments. Then, if in 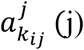 is a gap, we assume 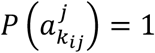. Then:

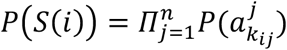
Step 6: Select candidates. The *i^th^* amino acid of protein P is selected as a candidate functional mutation if *P*(*S*(*i*)) is less than 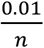, following the Bonferroni correction, and if 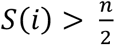. The method is schematically described in **Figure 5**

**Figure 5.**
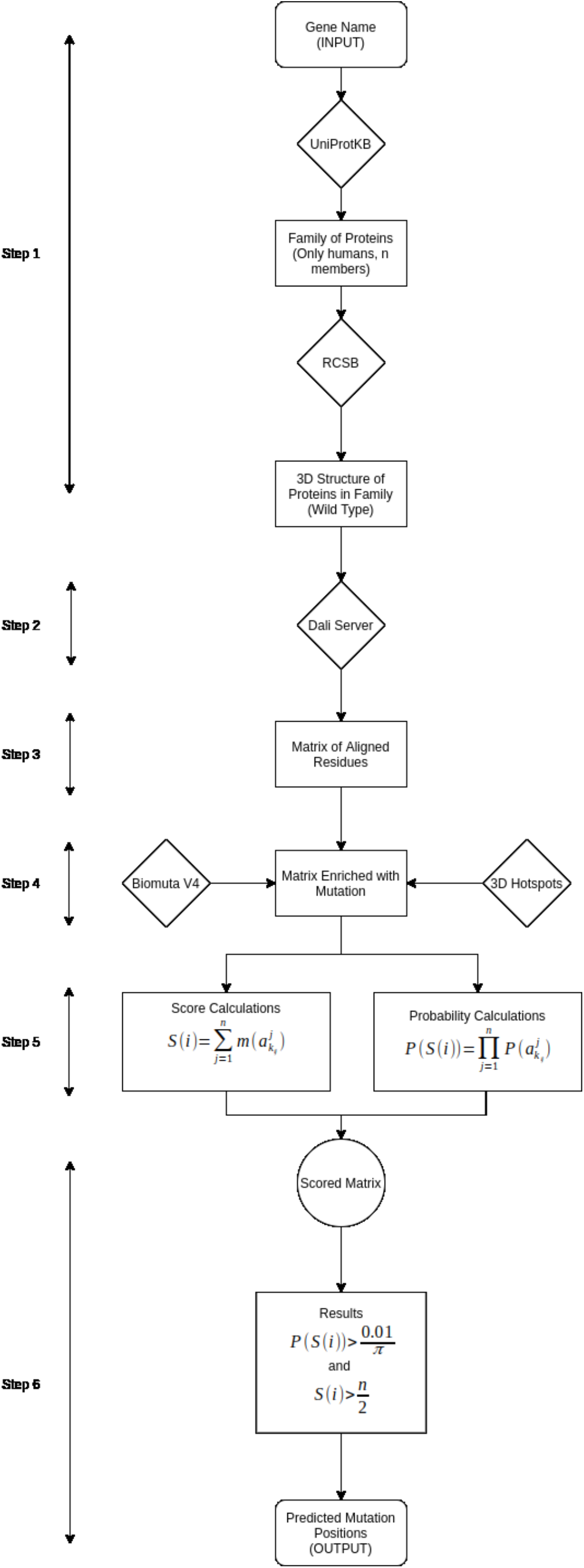
The PhiDsc workflow. The system begins by obtaining family members; the algorithm then obtains the 3D structures from RCSB; the algorithm aligns members pairwise with the input protein; mutations are then enriched in the alignments; finally, scores and probabilities are calculated.

### Leave-one-out cross-validation

In leave-one-out cross-validation (LOOCV), one data point from the training set remains excluded. For example, if there are n data points in the original sample, n-1 samples are used to train the model, and p points are used as the validation set. This is repeated for all combinations in which the original sample can be separated in this manner, and the error is averaged across all trials to calculate overall effectiveness. The number of possible combinations is equal to the original sample’s number of data points, or n.

*A_i_*(*P*) = {*P*_1_,*P*_2_,*P*_3_… *P*_n_. } – {*P*_i_} is considered as an input set for protein P, and the PhiDsc predictions for P are obtained by considering *A_i_*(*P*) as its protein family set. The set of predicted functional mutations is obtained for every 1 ≤ *i* ≤ *n*. A projected functional mutation is said to be robust if it is predicted across at least 80% of all rounds.

### Residue Interaction Network

RIN (Residue Interaction Network) is used to quantify the physical effect of the mutation on protein structure and function. In summary, Chang *et al.* demonstrated that if a mutation in a protein’s 3D structure is close to some hotspot mutations, the likelihood of this mutation being considered a hotspot mutation is high. The RINalyzer (49) module generates user-defined RINs from a 3D protein structure obtained from RCSB protein databank. RINerator considers different biochemical interaction types, such as contacts/clashes, hydrogen bonds, and hydrogen atoms and quantifies their individual strength as described in Chimera (50). RINalyzer is a Java plugin for Cytoscape(51), a free software platform for the analysis and visualization of molecular interaction networks. The results of interacting residues from RIN are compared to cBioPortal (11, 12) a dataset of mutations that are curated across cancer samples.

### Functional effect of candidate mutations on proteins

The effect of alterations in regions that were not identified as functional mutations experimentally can be calculated using a variety of methods. PhiDsc’s functional predictions are evaluated using six methods that, according to Stefl et al. (52), can be classified into three types:

The first group includes machine learning approaches that are trained on protein stability features and account for experimental conditions such as temperature, salt concentration, and pH values. Incorporating such parameters is critical for assessing the free-energy changes caused by mutations under near physiological conditions. This group includes I-Mutant2.0 (53) which uses SVM to estimate ΔΔG upon mutation, and PoPMuSiC-2.0 (54) which uses a mix of statistical potential and neural networks to estimate ΔΔG upon mutation.

The second group relies on evolutionary conservation data, with the assumption that changes at conserved positions in multiple sequence alignments are detrimental. Although these approaches do not directly predict the effect of mutations on protein stability, they are commonly used in conjunction with the methods mentioned above to achieve consensus predictions. This group includes SIFT (55), which uses sequence homology and site conservation to estimate the deleterious effect of mutations, and Provean (56), which predicts the functional impact of all types of protein sequence variations, including single amino acid substitutions, insertions, deletions, and multiple substitutions.

The third group uses structural information, assuming that a protein’s ability to function properly is determined by fundamental physicochemical properties that can only be derived from structures. This group includes CUPSAT(57), which estimates ΔΔG upon mutation using mean force atom pair and torsion angle potentials, and MutPred(58), which estimates detrimental effect of mutation using SIFT and gain/loss of structural or functional features predicted from sequences.

## DATA AVAILABILITY

This method is implemented in Python and the Source code and all tested data can be found on (https://github.com/hobzy987/PhiDSC-DALI). The software takes a UniProt Protein name as input and gives html file as output with aligned residues and probabilities, and a list of all residues sorted according to their score.

## ACKNOWLEDGEMENT

The authors thank Dr. Hossein Khiabanian for the insightful discssions and contribution provided for this work.

## Notes

### Competing Interest Statement

The authors have declared no competing interest.

